# Species richness increases fitness differences, but does not affect niche differences

**DOI:** 10.1101/823070

**Authors:** Jurg W. Spaak, Camille Carpentier, Frederik De Laender

## Abstract

A key question in ecology is what limits species richness. Modern coexistence theory presents the persistence of species as a balance between niche differences and fitness differences that favor and hamper coexistence, respectively. With most applications focusing on species pairs, however, we know little about if and how this balance changes with species richness. Here, we present the first mathematical proof that the average fitness difference among species increases with richness, while the average niche difference stays constant. Extensive simulations with more complex models and analyses of empirical data confirmed these mathematical results. Taken together, our work suggests that, as species accumulate in ecosystems, ever-increasing fitness differences will at some point exceed constant niche differences, limiting species richness. Our results contribute to the expansion of modern coexistence theory towards multi-species communities.

## Introduction

Explaining nature’s biodiversity is a key challenge for science (Hutchinson, 1957). One type of approach consists of focusing on the capacity of individual species to persist through time despite occasional pruning to low density or species interactions (Turelli, 1978). Modern coexistence theory is such an approach, and predicts species persistence in a community when niche differences overcome fitness differences. Niche differences measure the strength of negative frequency dependence, i.e. whether a focal species *i* can recover when reduced to low abundance (Adler *et al*., 2007; Chesson, 2000; Spaak & De Laender, 2020). Fitness differences measure the competitive ability of that species (Adler *et al*., 2007; Hart *et al*., 2018; Spaak & De Laender, 2020).

However, few applications of coexistence theory have explicitly focused on explaining biodiversity, i.e. the persistence of species in species-rich communities. Instead, most applications have considered two-species communities (Chesson, 2000; Letten *et al*., 2017), using a variety of approaches and case studies, including annual and perennial plants (Adler *et al*., 2018; Godoy & Levine, 2014), phytoplankton (Gallego *et al*., 2019; Narwani *et al*., 2013) and bacteria (Zhao *et al*., 2016), and under different environmental conditions (Bimler *et al*., 2018; Cardinaux *et al*., 2018; Grainger *et al*., 2019; Lanuza *et al*., 2018; Matías *et al*., 2018; Napier *et al*., 2016; Wainwright *et al*., 2019). To our knowledge only three empirical studies that report niche and fitness differences in communities composed of more than two species (hereafter multi-species communities) (Chu & Adler, 2015; Petry *et al*., 2018; Veresoglou *et al*., 2018). However, none of these three studies explain how niche and fitness differences change with species richness. In order to understand how niche and fitness differences co-determine species persistence in multi-species communities, we need to understand how both variables change when adding species to a community.

Multi-species communities possess at least four complexities that are absent from two-species communities, which may affect niche and fitness differences, and therefore how we interpret coexistence in multispecies communities. (1) First, a multi-species community can host more interaction types than species pairs, e.g. competitive, mutualistic or trophic interactions. Species richness increases the number of possible interactions and the number of combinations of these interaction types. Several metrics exist to summarize this diversity of interaction types and study their implications for community dynamics (Fontaine *et al*., 2011). (2) Second, two-species communities are always fully connected and correlations between interspecific interactions (Barabás *et al*., 2016b) become irrelevant since there is only a single pair of interspecific interactions. In contrast, in an *n*-species community there may be anywhere from *n* − 1 to 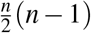 connections, and the interspecific effects of species *j* on species *i* can be positively or negatively correlated with the interspecific effects *i* has on *j* (Barabás *et al*., 2016b). May (1972) and Allesina & Tang (2012, 2015) and have shown that connectance and correlation can have large effects on the local stability of multi-species communities. We therefore expect these factors to influence coexistence as well. (3) Third, higher-order interactions, through which a third species changes the interaction between a species pair, are by definition absent from two-species communities (but see Letten & Stouffer (2019); Levine *et al*. (2017)). Such higher-order interactions have been found empirically, for example, in communities composed of phytoplankton, bacteria, and ciliates (Mickalide & Kuehn, 2019). In that study, bacteria coexisted with phytoplankton and ciliates, but all three functional groups did not coexist. In the three-species community, phytoplankton inhibited bacterial aggregation, leaving the latter more vulnerable to predation by ciliates. (4) Fourth, indirect effects, whereby a third species changes the dynamics of a species pair by directly interacting with both partners at the same time, are by definition absent from two-species communities (Walsh, 2013). We will refer to these four complexities throughout the text with (1) interaction types, (2) interaction matrix structure, (3) higher-order interactions and (4) indirect interactions.

Studying multi-species coexistence is challenging both theoretically and experimentally. Theoretically speaking, the methods to analyse coexistence via niche and fitness differences in a multi-species community were not available until recently (Carmel *et al*., 2017; Carroll *et al*., 2011; Saavedra *et al*., 2017; Spaak & De Laender, 2020). Experimentally speaking, studying coexistence of multiple species is resource-demanding. For instance, in the simple case of linear direct interactions among species (i.e. as in Lotka-Volterra models) the number of experiments needed to parametrize the community is quadratic in species richness (but see Maynard *et al*. (2019)). Considering higher-order interactions will consequently result in an even higher experimental load. For example, measuring higher-order interactions, sensu Letten & Stouffer (2019), would require 39 experiments in a three species community.

In this paper we investigate the balance between niche and fitness differences along a gradient of species richness. More specifically, we ask how niche and fitness differences change as the number of species increases in randomly assembled communities, and how the additional complexities (1)-(4) influence these changes. We do so using four independent methods that rely on a novel definition for niche and fitness differences that is able to analyse multi-species coexistence through niche and fitness differences (Spaak & De Laender, 2020), and a new compilation of species interaction data. First, we derive equations that quantify how niche and fitness differences respond to species richness in a community with linear interactions and simple cases of higher-order interactions. Second, we give an intuitive explanation of these responses based on the Mac-Arthur consumer-resource model. Third, we perform simulations with more complex models. We run these simulations as a full-factorial virtual experiment, varying direct interactions (type, correlation, connectance), higher-order interactions and indirect interactions. Fourth, we compile data from the literature on empirically measured species interaction matrices and compute niche and fitness differences as a response to species richness. We then compare our results obtained via random species assembly to a community with non-random assembly. All methods support the same general conclusion: species richness does not affect niche differences, but increases fitness differences. Importantly, these conclusions are independent of the four complexities (1)-(4).

## Methods

### Model assumptions

To include the various additional complexities of multispecies communities we use a generalized Lotka-Voltera model with *n* species containing second-order interactions to model the per-capita growth rates *f*_*i*_ (*N*_*i*_, ***N***^*−i*^):

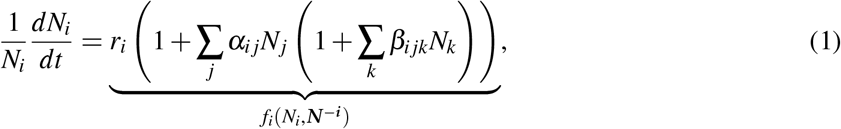

where *N*_*i*_ is the density of the focal species *i*, **N**^−*i*^ is the density of the resident community (vector of length *n* − 1), *r*_*i*_ is its intrinsic growth rate, and *α*_*ij*_ and *β*_*ijk*_ are first (or linear) and second-order species interactions, respectively. A positive *α*_*ij*_ indicates a positive interaction between species *i* and *j* (facilitation). Negative *α*_*ij*_ on the other hand indicate negative interactions (competition). When *β*_*ijk*_ is positive or negative species *k* will intensify or weaken the relationship between species *i* and *j*, respectively (second-order interaction). The inclusion of third order interactions did not affect any of our results. Throughout the manuscript we assume *α*_*ii*_ *=* −1 with no loss of generality as it can be achieved by rescaling of empirical data. Furthermore, for most of the manuscript we assume a random community assembly. That is *α*_*ij*_ and *α*_*ik*_ are sampled from any independent distributions for *k* ≠ *j* and similarly for *β*_*i j k*_.

### Niche and fitness differences

There exist a total of eleven definitions for niche and fitness differences with various assumptions about the community model (Spaak & De Laender, 2020). Five of these definitions can be applied to multi-species communities (Carmel *et al*., 2017; Carroll *et al*., 2011; Chesson, 2003; Saavedra *et al*., 2017; Spaak & De Laender, 2020). Optimally, we would compute niche and fitness differences according to all five methods to understand how species richness affects coexistence.

However, this is not possible, as four of the five community models do not readily apply to the communities investigated here. The definitions of Carmel *et al*. (2017) assume that the species have a minimal growth rate, the mortality rate, it therefore best applies to resource competition models, but it does not apply to Lotka-Volterra models. Chesson (2003) was developed for environmental or spatial fluctuations, which we do not consider here. Furthermore, Chesson (2003) works best for communities driven by only one limiting factor (Barabás *et al*., 2018), the community models investigated here, however, have as many limiting factors as they have species. For such communities the scaling factors are not uniquely defined and the results will therefore depend on the non-unique choice of the scaling factors (Barabás *et al*., 2018; Song *et al*., 2019). Saavedra *et al*. (2017) can only be applied to communities without higher order interactions. Cenci & Saavedra (2018) propose and extension to include non-linear per-capita growth rates, however, the interaction matrix must still be linear, which is not the case for the communities investigated here. Carroll *et al*. (2011) can only be applied to communities without strong inter-specific facilitation, that is the intrinsic growth rate must be higher than the invasion growth rate. This excludes about two-thirds of our theoretical and empirical data. One could ask how species richness affects niche and fitness differences according to these last two definitions if the community model were restricted to competitive Lotka-Volterra communities, yet, there is an additional problem. Niche and fitness differences as computed by these two definitions allow inference of coexistence only in two-species communities, not in multi-species communities (Spaak & De Laender, 2020). That is, two different multi-species communities may have identical niche and fitness differences but different outcomes of coexistence (e.g. all species persist in one community, but not in the other). Since we ask how niche and fitness differences jointly determine coexistence in multi-species communities, these methods are not suited to gain new insight. Consequently, of the eleven available definitions of niche and fitness differences only the definition of Spaak & De Laender (2020) applies to the community models and the research question investigated here.

Spaak & De Laender (2020) base their definition of niche and fitness differences (𝒩_*i*_ and ℱ_*i*_) on the comparison of species growth rates in various conditions. If the two species *(i* and *j)* of a two-species community have completely separated niches (𝒩_*i*_ = 𝒩_*j*_ *=* 1), the species *i* will grow in presence of *j* as it would in its absence, which implies 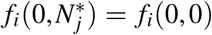 where 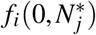 is the per capita growth rate of species *i* when it invades a community with *j* at equilibrium density 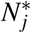, known as the invasion growth rate (Carroll *et al*., 2011; Chesson, 1994; Ellner *et al*., 2019) and *f*_*i*_(0,0) is the intrinsic growth rate of species *i*. Conversely, if the two species have exactly the same niche (𝒩_*i*_ = 𝒩_*j*_ *=* 0) they have equivalent effects on each other. It then holds that 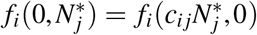 where *c*_*ij*_ is the conversion factor allowing to express individuals of species *j* as individuals of species *i*.

Unfortunately, the *c*_*ij*_ are defined implicitly as the solution of an equation, which often does not have a closed form solution, but must be computed numerically. However, intuitively the *c*_*ij*_ are simple. The conversion factors *c*_*ij*_ change the frequency of species *i* and *j*, but they do not affect the total density. That is, two communities with densities 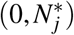 and 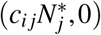 have the same total density, but different frequencies of species *i*. In the first community species *i* has 0 frequency, while in the second it has 100% frequency, as 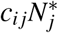 is a density of species *i*. We refer to Spaak & De Laender (2020) and the appendix S1 for a mathematical definition of *c*_*ij*_.

Further intuitive insight can be gained from a Lotka-Volterra model in which the *c*_*ij*_ can be computed explicitly as 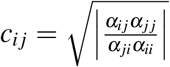 (Appendix S2). Note that the arrangement of the interaction coefficients *α*_*ij*_ differs from both, the niche and the fitness differences. The numerator (*α*_*ij*_ *α*_*jj*_) is the competitive effect of species *j*, while the denominator *(α*_*ji*_*α*_*ii*_) is the competitive effect os species *i*. The *c*_*ij*_ therefore keep total density constant by scaling the effects of the two species.

Niche differences are then defined via interpolation between these two extreme cases:

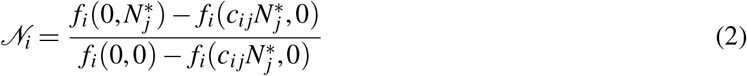

This equation of niche differences can be interpreted as the relative frequency dependence of species *i*. The numerator assesses frequency dependence, as it compares the growth rate of species *i* when it has 0% frequency to when it has 100% frequency, notably at the same converted density. The denominator assesses density dependence, as it compares the growth rate of species *i* at 0 density to when it has non-zero density. Importantly, in both growth rates species *i* has 100% frequency. This definition maps positive frequency dependence, negative frequency dependence and facilitation to 𝒩_*i*_ < 0,0 < 𝒩_*i*_ < 1, and 1 < 𝒩_*i*_ respectively.

Similarly, they define fitness differences as the scaled growth rate in the absence of niche differences

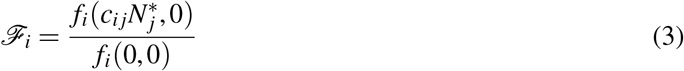

Zero fitness differences imply that the species have equal competitive strength, as 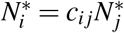 that is species will have the same equilibrium density after conversion with *c*_*ij*_. Competitive subordinate species then have negative fitness differences, as 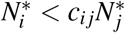, conversely competitive dominant species have positive fitness differences. Note that ℱ_*i*_ *=* 0 means no fitness differences and more negative ℱ_*i*_ means stronger fitness differences. Importantly, these definitions of niche and fitness differences agree with the original definitions on the two species Lotka-Volterra models (Chesson, 2013; Chesson & Kuang, 2008), they are therefore not newly defined niche and fitness differences, but rather a generalization thereof (Appendix S1, Spaak & De Laender (2020)).

Additionally, this definition very naturally generalizes to multi-species communities. The three growth rates remain conceptually the same. The intrinsic growth rate *f*_*i*_ (0,**0**), where **0** denotes the absence of all other species. The invasion growth rate *f*_*i*_ (0,**N**^(−*i*,*)^), where **N**^(−*i*,*)^ denotes the equilibrium density of the community in absence of species *i*. And the no-niche growth rate 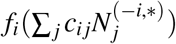, where all densities of the resident species have been converted to densities of the focal species. Doing so makes clear that 𝒩_*i*_ and ℱ_*i*_ are species-specific properties, i.e. in general we have 𝒩_*i*_ ≠ 𝒩_*j*_ and ℱ_*i*_ ≠ ℱ_*j*_ in multi-species communities. However, 𝒩_*i*_ and ℱ_*i*_ compare the effect of the focal species on itself to the effect of the rest of the community on the focal species. They therefore do depend on the traits of the other species in the community.

### Analyses and Simulations

We first examined analytically how 𝒩_*i*_ and ℱ_*i*_ change with species richness. We found a generic solution for first-order interactions and for a simplified case of higher-order interactions.

Second, we designed a full-factorial virtual experiment in which we numerically computed 𝒩_*i*_ and ℱ_*i*_ for a wider range of different simulated communities (see table 1). For these we solve numerically for equilibrium densities and invasion growth rates using the’fsolve’ function from the scipy package in Python. Communities with higher order interactions can have multiple equilibria (Aladwani & Saavedra, 2019). To compare communities with and without higher order interactions we computed 𝒩_*i*_ and ℱ_*i*_ only for one equilibrium in communities with higher order interactions. Specifically, we always chose the equilibrium that is closest to the equilibrium of the community without higher order interactions. The factors were (i) first-order interaction type (competitive, facilitative or both, i.e. *α*_*ij*_ < 0, > 0 or unrestricted); (ii) first-order interaction strength (strong or weak); (iii) connectance of the interspecific interaction 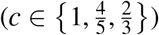; (iv) correlation between the interspecific interaction (*ρ*(*α*_*ij*_, *α*_*ji*_) = *ρ*_*ij*_(*β*_*ijk*_,*β*_*jik*_) ∈ {− 1,0,1}); (v) inclusion of indirect effects (present or absent); (vi) second-order interaction type (*β*_*ijk*_ < 0, > 0, unrestricted, or absent); To exclude indirect effects we set equilibrium densities of resident species to their monoculture equilibrium density. In this way, we cancel out interactions among residents that will change the residents’ densities. The intrinsic growth rate *r*_*i*_ does not affect 𝒩_*i*_ and ℱ_*i*_, therefore we set it to 1 in all simulations.

**Table 1:** Design of full factorial virtual experiment.

| Factor | Parameter | Levels | Interpretation | Complexity investigated |
| --- | --- | --- | --- | --- |
| Interaction type<br>first-order | $\alpha_{ij}$ | $< 0$<br>$> 0$<br>no restriction | competition<br>facilitation<br>mixed | (1) |
| Strength of interaction<br>first-order | $\alpha_{ij}$ | strong<br>weak | | (1) |
| Connectance | $P(\alpha_{ij} \neq 0)$ | $1, \frac{4}{5}, \frac{2}{3}$ | | (2) |
| Interaction<br>correlation | $\text{cor}(\alpha_{ij}, \alpha_{ji})$<br>$\text{cor}_{ij}(\beta_{ijk}\beta_{jik})$ | 1<br>0<br>-1 | equal<br>unrelated<br>opposite | (2) |
| Presence of<br>indirect effects |  | Yes<br>No | absent<br>present | (4) |
| Interaction type<br>second-order | $\beta_{ijk}$ | $> 0$<br>$< 0$<br>no restriction<br>$= 0$ | intensify<br>weaken<br>mixed<br>no second-order | (1) and (3) |

This design leads to a total of 3 · 2 · 3 · 3 · 2 · 4 = 432 parameter settings. We ran 1000 repetitions for each of the five species richness levels (2 ≤ *n* ≤ 6), leading to a total of 432 · 5 · 1000 = 2′160′000 simulations. We parametrized the first-order interactions using distributions of empirically obtained first-order interactions (see supplementary informations S2). We sampled “strong” first-order interactions between the *Q*_*1*_ − 1.5 *(Q*_*3*_ − *Q*_*1*_) and *Q*_*3*_ + 1.5 *(Q*_*3*_ − *Q*_*1*_) of those distributions, where *Q*_*1*_ and *Q*_*3*_ are the first and third quartile, to remove outliers (defined as species interactions outside this range, Appendix S3). Similarly, we sampled “weak” first-order interactions between the *Q*_1_ and *Q*_3_ of the empirical distributions of interaction strength. We fitted linear regressions to measure the effect of species richness on 𝒩_*i*_ and ℱ_*i*_. With this approach we were able to investigate the effects of all complexities (1)-(4).

### Literature data

We found three review papers that collected multi-species Lotka-Volterra interaction matrices (Adler *et al*., 2018; Fort, 2018; Keddy & Shipley, 1989), representing a total of 33 interaction matrices, ranging from 3 to 9 species, and containing 29 plant, 2 phytoplankton, 1 zooplankton and 1 ciliate communities. We normalized all these data such that *α*_*ii*_ *=* −1. The interaction matrices were obtained through pairwise experiments, measuring the interspecific effect of one species on the other. For each multi-species community we constructed all possible sub-communities with at least two species, leading to a total of 2544 communities that varied in species richness from 2 to 9. We excluded all communities in which not all interaction strengths were available, e.g. because of a “NA” entry in the sampled sub-community, leading to 2296 communities. For 1376 communities we could not compute 𝒩_*i*_ and ℱ_*i*_. That is because, like any method seeking to quantify frequency dependence, our approach is based on invasion analysis: the capacity of an invader to grow with the other species at their non-zero equilibrium. Thus, one must be able to compute the invasion growth rate of each species, which is the per capita growth rate 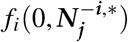 when the focal species *i* is nearly absent (mathematically equal to 0) and the other species are at their equilibrium density ***N***^*−i\**^. 𝒩_*i*_ and ℱ_*i*_ are thus only computable for communities where all species in each subcommunity (the community without the invading species) coexist stably. We therefore computed 𝒩_*i*_ and ℱ_*i*_ for the remaining 920 communities (see Appendix S4).

For each interaction matrix obtained from the literature we computed 𝒩_*i*_ and ℱ_*i*_ using equation 2 and 3. For each of the 33 interaction matrices, we regressed 𝒩_*i*_ against species richness of the sub-communities. These data contained many outliers, which skewed the results of our linear regressions. We therefore used a Theil-Sen estimator for the slope, which is more robust to outliers than linear regression based on least squares (Sen, 1968). We fitted (using least squares) a saturating function 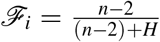 for the fitness differences. This saturating response was chosen for ℱ_*i*_, because our analytical results suggested such a response.

## Results

### Analytical solutions

For the linear Lotka-Volterra model without higher-order interactions (i.e. *β*_*ijk*_ *=* 0), we can explicitly compute ℱ_*i*_ and 𝒩_*i*_ the fitness and niche differences of species *i* in the multispecies community (see supplementary informations S1):

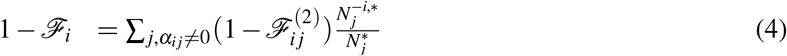

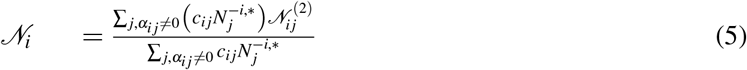

with 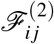 and 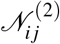 the fitness and niche difference of species *i* in a two-species community consisting of species *i* and 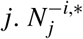 is the equilibrium density of species *j* in the absence of species *i*. 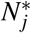 is the equilibrium density of species *j* in monoculture and *c*_*ij*_ is the conversion factor from species *j* to species *i* (see methods). The sums are taken across all species *j* with which species *i* interacts directly, i.e. *α*_*ij*_ ≠0.

Eq. 4 and 5 illustrate our two main results. First, 1 − ℱ_*i*_ is the weighted *sum* of the two-species fitness differences 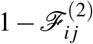, across all species pairs with which *i* interacts. The weights 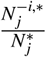 are the relative yields, as known from biodiversity ecosystem functioning research (Fox, 2005; Hector *et al*., 2001). The effect of species richness on fitness differences will therefore be similar to the effect of species richness on the sum of the relative yield, known as the relative yield total 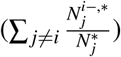, which is known to increase with species richness in many communities (Carroll *et al*., 2011; Grace *et al*., 2016; Loreau, 2004). Hence,1 − ℱ_*i*_ (and therefore of ℱ_*i*_ decreases) will on average increase with species richness. Second, 𝒩_*i*_ is the weighted *average* of the two-species niche overlaps 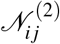. Hence, species richness will, on average, not affect 𝒩_*i*_ Since we did not make assumptions about the *α*_*ij*_, these results are independent of the details of interspecific interactions, i.e. the results apply regardless of complexities (1) and (2).

Equation 5 and 4 can be approximated by assuming constant interspecific interaction strength 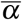 (see supplementary informations 1). This yields 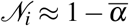 and 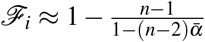, from which it is clear that 𝒩_*i*_ is independent of species richness *n* and ℱ_*i*_ is an increasing but saturating function of species richness.

The saturation occurs because the sum of the weights 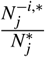, the relative yield total, will saturate as well in the Lotka-Volterra model (Loreau, 2004; Spaak *et al*., 2017). Including other complexities does not affect these two main results (Appendix S2).

### Link to resource competition

The fact that 1 − ℱ_*i*_ is a weighted sum, while 𝒩_*i*_ is a weighted average makes intuitive sense when realising that the interaction coefficients *α*_*ij*_ can, under certain conditions, be related to resource utilisation (Chesson, 1990; MacArthur, 1970). We want to compute the 𝒩_1_ and ℱ_1_ of the yellow focal species (yellow, Fig. 1) in presence of one up to five competitors (Fig 1 A-E). We assume that the species only differ in resource utilisation rates, not in other parameters such as mortality.

**Figure 1:**
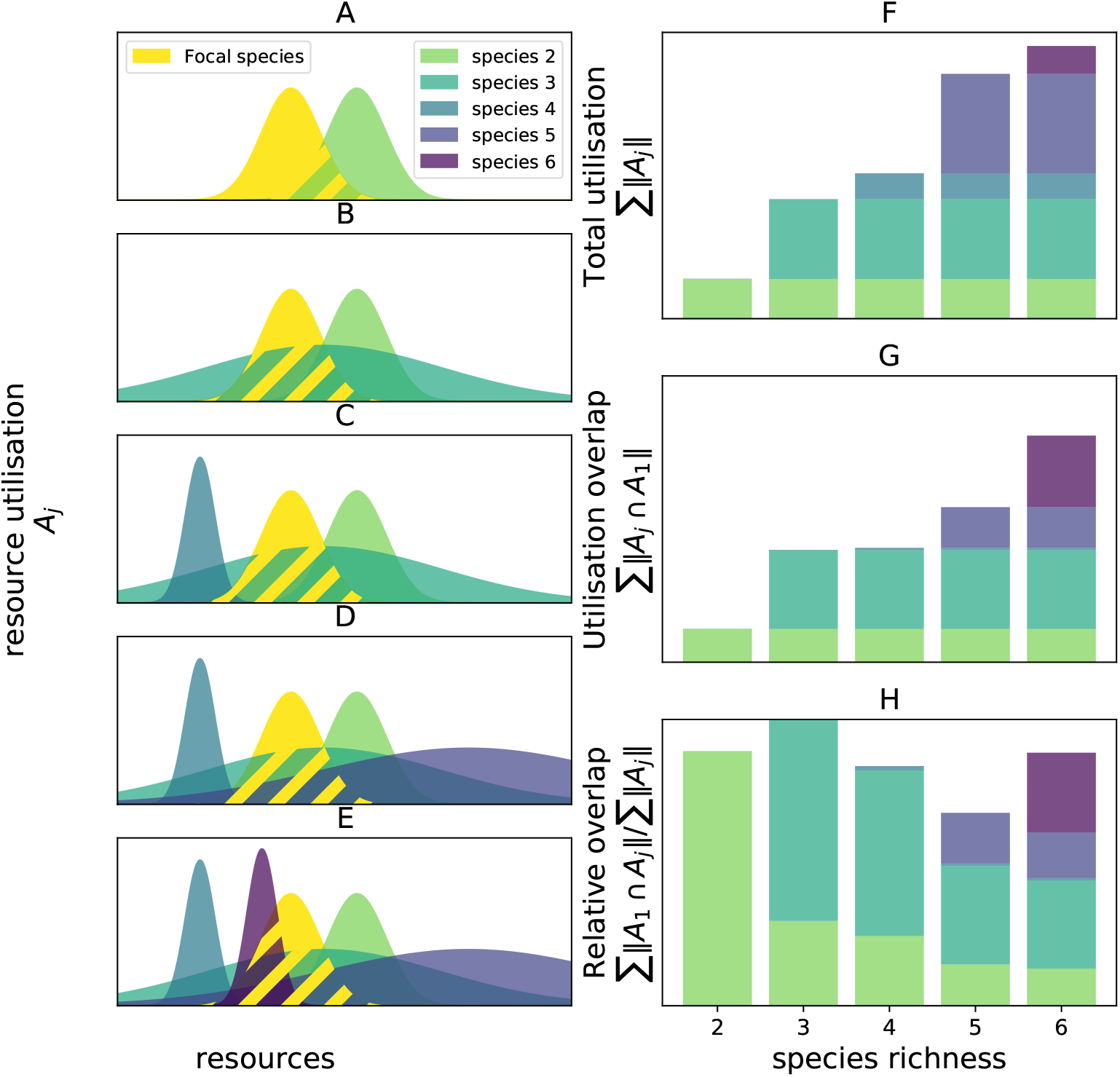
A-E: Resource utilisation of the yellow focal species and its competitors. F: Increasing the species richness will increase the total utilisation of the resident species ∑ _*j*_ ‖*A*_*j*_‖. Similarly, we expect 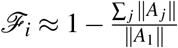 to increase with species richness, as it scales with the ratio of total resource consumption. G: The amount of shared resources (hatched region from panels A-E) increases with species richness. H: As both the amount of shared resources increase (panel G) and the total utilisation (panel F) increase, we expect the ratio to be independent of species richness. Similarly, we expect 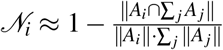 to be, on average, independent of species richness. F-H: The colours of the bar correspond to the contribution of each of the resident species to the total of the bar.

In a two species community, the species with the higher total utilisation rate, denoted ‖*A*_*i*_‖ the area under the curve *A*_*i*_, will have a competitive advantage and consequently the higher fitness difference. One could therefore intuit that the fitness difference is linked to the ratio of total resource utilisation rates, i.e. 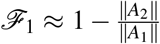. Fitness differences therefore increase with species richness, as each competitor increases the total resource utilisation rates of all competitors combined (Fig. 1 F), i.e.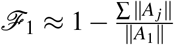. It turns out that this intuition is almost correct; we only have to add weights to the sum according to the densities of the species at equilibrium (compare this equation to eq. 4). ℱ_*1*_ thus increases (becomes more negative), as species richness increases (note that ℱ_*i*_ *=* 0 means no fitness differences and more negative ℱ_*i*_ means stronger fitness differences).

Intuitively, 1 − 𝒩_*i*_ is the proportion of shared resources between the focal species and its competitors, that is the amount of shared resources scaled by the total consumption of the species. In a two species community we therefore intuit 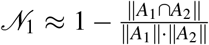. Increasing species richness will increase the sum of shared resources, as the focal species will share resources with each competitor (Fig. 1 G), but also the sum of the total consumption of the species (Fig. 1 F). We therefore expect 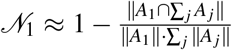 to be independent of species richness (Fig 1 H). Again, this intuition is correct up to the inclusion of the weights according to the species equilibrium densities.

### Full-factorial simulations

The simulations with strong first-order interactions only partially seem to confirm the predictions made by theory (Fig. 2). That is, 𝒩_*i*_ is not invariant but approaches 1 as species richness increases (Fig. 2 A), which seems to contradict the theoretical results. Yet, species richness does not directly affect 𝒩_*i*_, but rather affects the average interaction strength 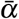, which in turn affects 𝒩_*i*_, which can be approximated by 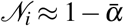 (see Appendix S2). That is because, by design, any method based on invasion growth rates (such as those to compute 𝒩_*i*_ and ℱ_*i*_) can only be applied to communities in which invasion analysis is possible. Hence, too strong negative interactions prevent the invasion into highly-diverse communities, and will often impede feasible *n −* 1 subcommunities to begin with (Kokkoris *et al*., 2002). Hence, species richness selects against communities with overly strong negative interactions, which leads to on average weaker interactions at higher species richness (see supplementary Fig. S3). Similarly, species densities in communities with strong positive interactions will tend to grow to infinity, and more so in species-rich communities, because interspecific facilitation is stronger than intraspecific limitation (self-regulation). Again, species richness selects against strong positive interactions, weakening the average interaction strength (see supplementary Fig. S3). This selection of weak (negative and positive) interspecific interactions causes 𝒩_*i*_ to approach 1. ℱ_*i*_ increases with species richness for all parameter settings, as predicted by the theory (see Fig 2 B).

**Figure 2:**
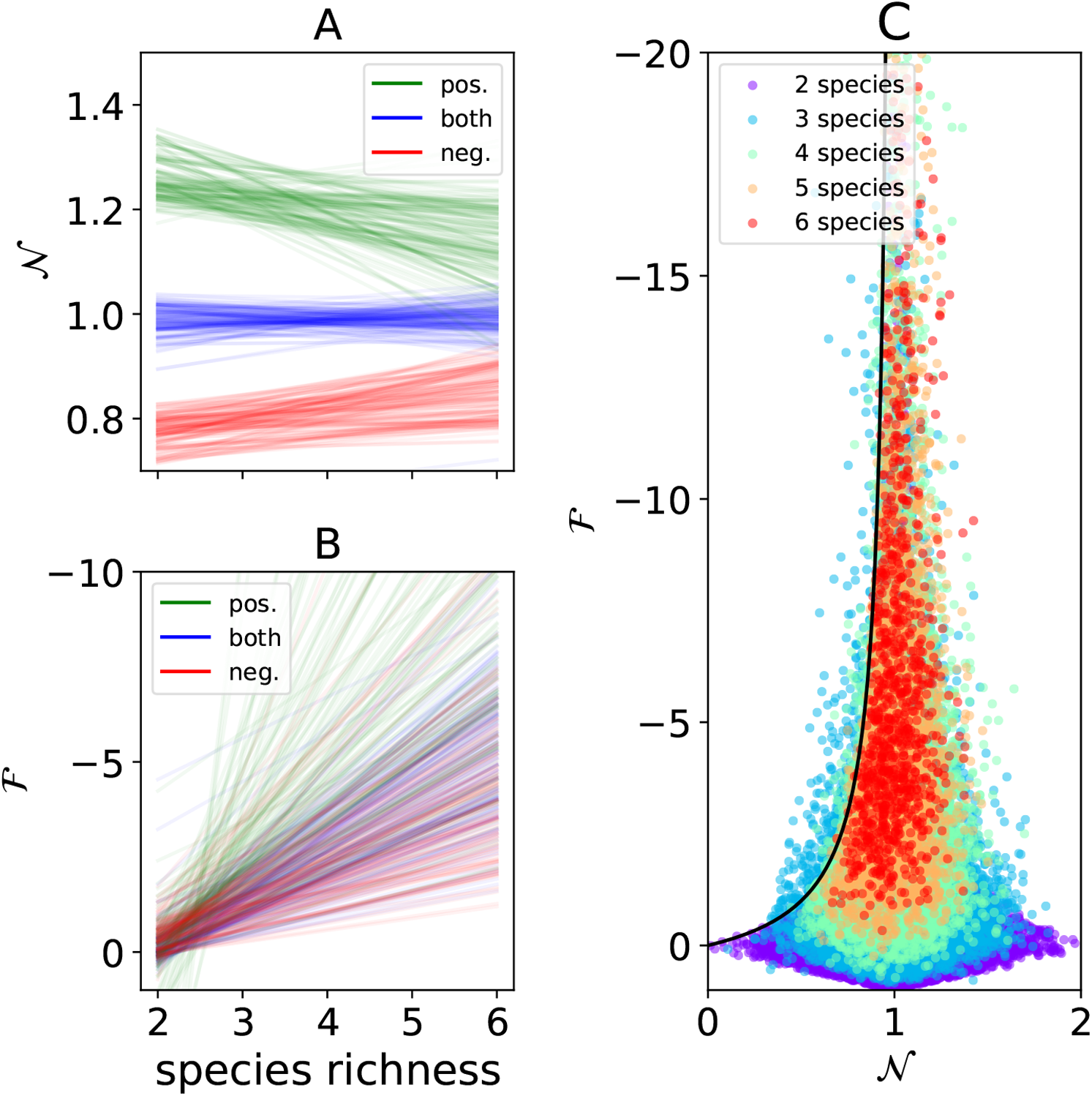
𝒩_*i*_ and ℱ_*i*_ as a function of species richness in simulated communities with strong first-order interspecific interactions. A: Contrary to predictions from theory, niche differences change with species richness when first-order interspecific interactions are either positive (green) or negative (red; results for unconstrained interspecific interactions are shown in blue). However, this is because interaction strength decreases with increasing species richness in these cases (see Fig. S2). Each line represents a linear regression of niche differences as a function of species richness for one factorial setting of the full-factorial experiment (see table 1). B: Species richness, however, makes fitness differences more negative (i.e. larger). Note the differences in y-scale between panel A and B. C: Distribution of 𝒩_*i*_ and ℱ_*i*_ for simulated theoretical communities that are fully connected, and exhibit first-order interactions without correlations, i.e. similar to the experimental communities (see Fig. 4). Each dot represents 𝒩_*i*_ and ℱ_*i*_ of one species in a community composed of 2-6 species (see colour legend). The black line indicates the persistence line, species below this line are assumed to persist in the community. Note the inverted y-axis.

The simulation results based on weak interaction strengths allow assessing the direct effect of species richness on 𝒩_*i*_ and ℱ_*i*_ without the confounding effect of species richness on interspecific interaction strength *α*_*ij*_. In these simulations, the effect of species richness on interspecific interactions was much weaker (see supplementary Fig. S3). These simulations confirmed our theoretical results; 𝒩_*i*_ was on average unaffected by species richness (see Fig. 3 A) and ℱ_*i*_ increased with species richness (Fig. 3B). We illustrate how 𝒩_*i*_ and ℱ_*i*_ values jointly varied with species richness, using weak interaction strength: no higher-order interactions *(β*_*ijk*_ *=* 0), no correlation between the *α*_*ij*_, and maximum connectance (Fig. 3 C). Again, these results hold independently of the complexities (Appendix S3).

**Figure 3:**
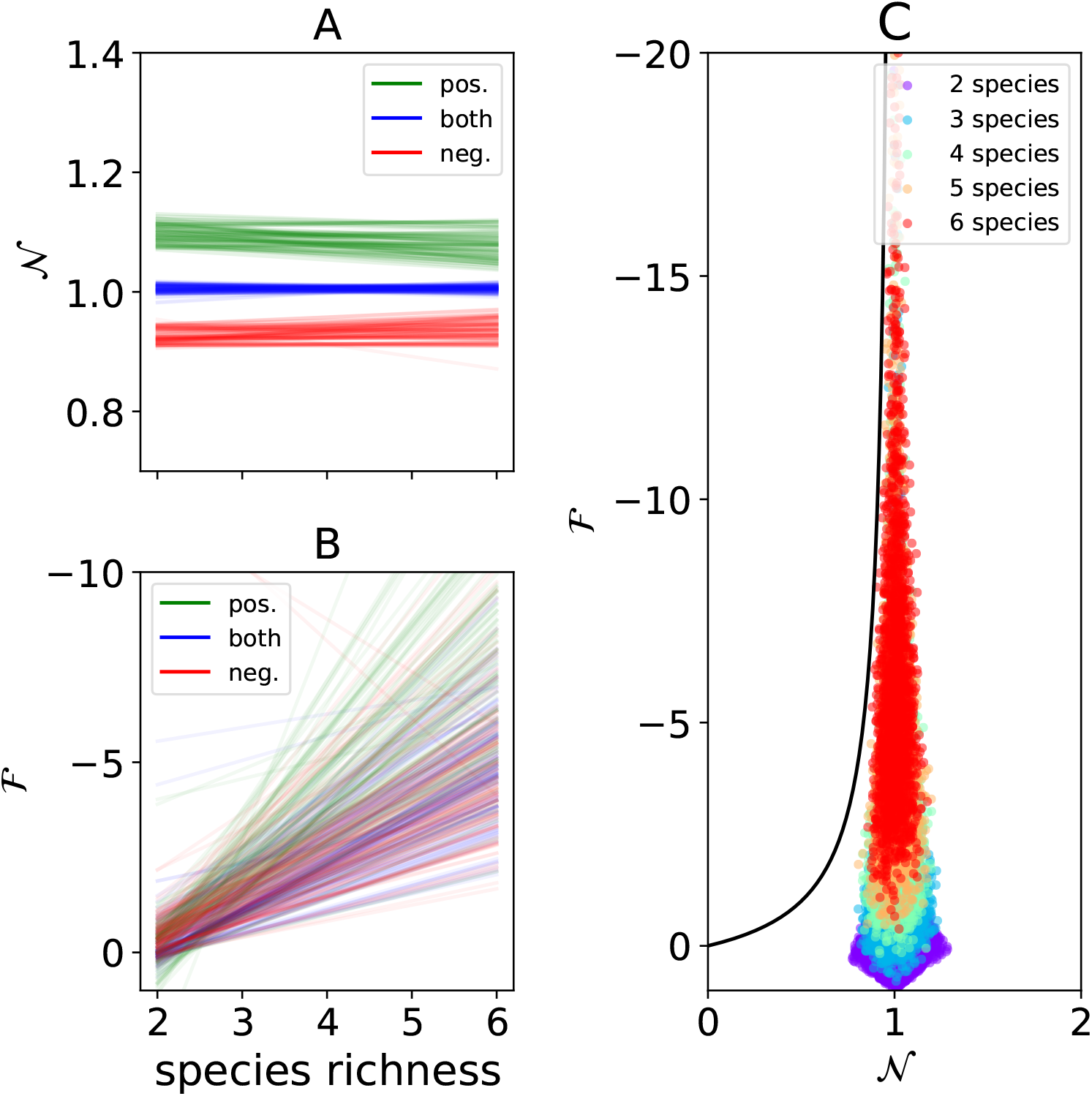
As Fig. 2, but with weak first-order interspecific interactions. A: As predicted by the mathematical results, species richness does not affect niche differences, because communities with different species richness had comparable interaction strengths. B: Species richness, however, makes fitness differences more negative (i.e. larger). C: Distribution of 𝒩_*i*_ and ℱ_*i*_ for simulated theoretical communities that are fully connected, and exhibit first-order interactions without correlations, i.e. similar to the experimental communities (see Fig. 4).

### Literature Data

The results for the empirical communities reflect those obtained for the simulated communities. The absolute values of the slopes of the linear regressions of 𝒩_*i*_ were small (< 0.05) for all but 6 datasets. The slope for the overall regression of 𝒩_*i*_ against species richness (Fig. 4A, blackline) was small (−0.028). ℱ_*i*_ increased with richness in all but one dataset. Overall, we conclude that the response of 𝒩_*i*_ and ℱ_*i*_ to richness for empirical communities did not qualitatively differ from that of randomly generated communities.

**Figure 4:**
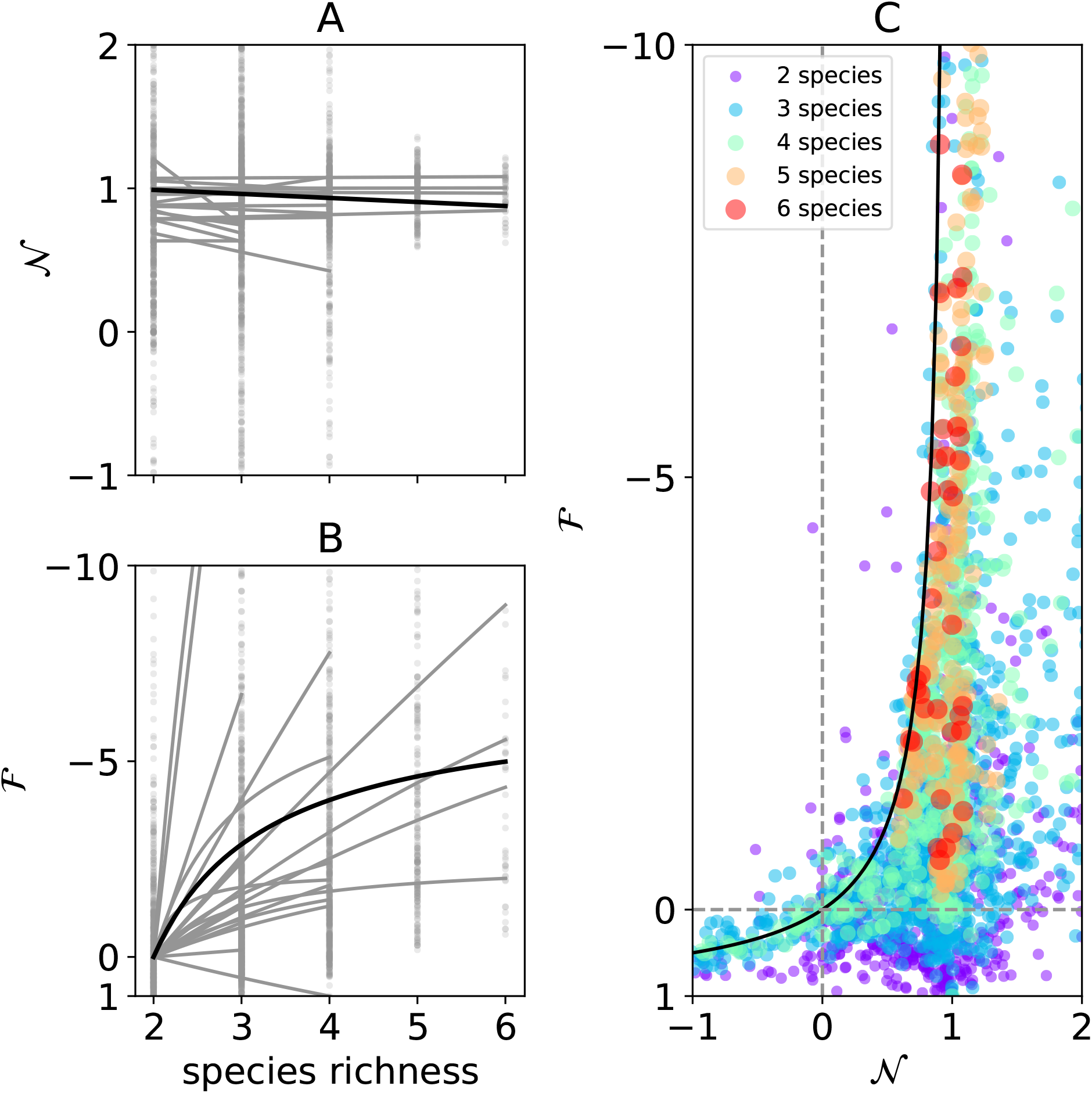
Similar to Fig 2 for empirically observed communities. Each grey line corresponds to a fit of a linear (𝒩_*i*_, A) and saturating (ℱ_*i*_, B) regression model to one dataset. The black line represents a fit through all 𝒩_*i*_ (A) and ℱ_*i*_ (B) values. Grey dots in panel A and B represent the raw 𝒩_*i*_ and ℱ_*i*_ values. Facilitation, i.e. species having a positive net effect on another, and therefore 𝒩_*i*_ > 1 is common in the datasets we found. We highlight one specific three-species community (grey line) where all species coexist, even though species *a* has 𝒩_*i*_ < 0, a property associated with priority effects and therefore exclusion. Axis from C are truncated to show ∼95% of all data points.

The empirical data also revealed cases in which coexistence is possible even though some of the species have negative 𝒩_*i*_. This is possible as long as ℱ_*i*_ is sufficiently positive such that 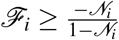. A total of 95 (4.1%) communities were found with species persisting despite having negative 𝒩_*i*_.

### Non-random community assembly

So far we have focused on a random community assembly, where the location and width of the resource utilisation *A*_*i*_ of species *i* were chosen randomly (Fig. 1 A-E). We here focus on two different possibilities of non-random community assembly. First, given a species pool, we can rearrange the order of species arrival. By choosing a non-random community assembly, species richness can increase or decrease niche differences, but will always increase fitness differences (Appendix S5). Additionally, averaged over all possible community assemblages species richness does not affect niche differences.

Alternatively, one might ask how species richness affects niche and fitness differences, if species optimize their resource utilisation according to the prevailing species richness (Barabás *et al*. (2016a), Fig. 5 A-E, Appendix S6). Importantly, in this case the two-species community is *not* a subcommunity of the three-species community. This links to the traditional question of species packing, which is how close can species be packed in a given environment and still coexist (Barabás *et al*., 2014; MacArthur, 1970). In this scenario, species richness still increases fitness differences (Fig. 5 F). However, because of the limited niche space, species in species rich communities are located closer to each other and therefore have a stronger interaction strength on average (Fig. 5 G). Niche differences therefore slightly decrease with species richness (Fig. 5 H).

**Figure 5:**
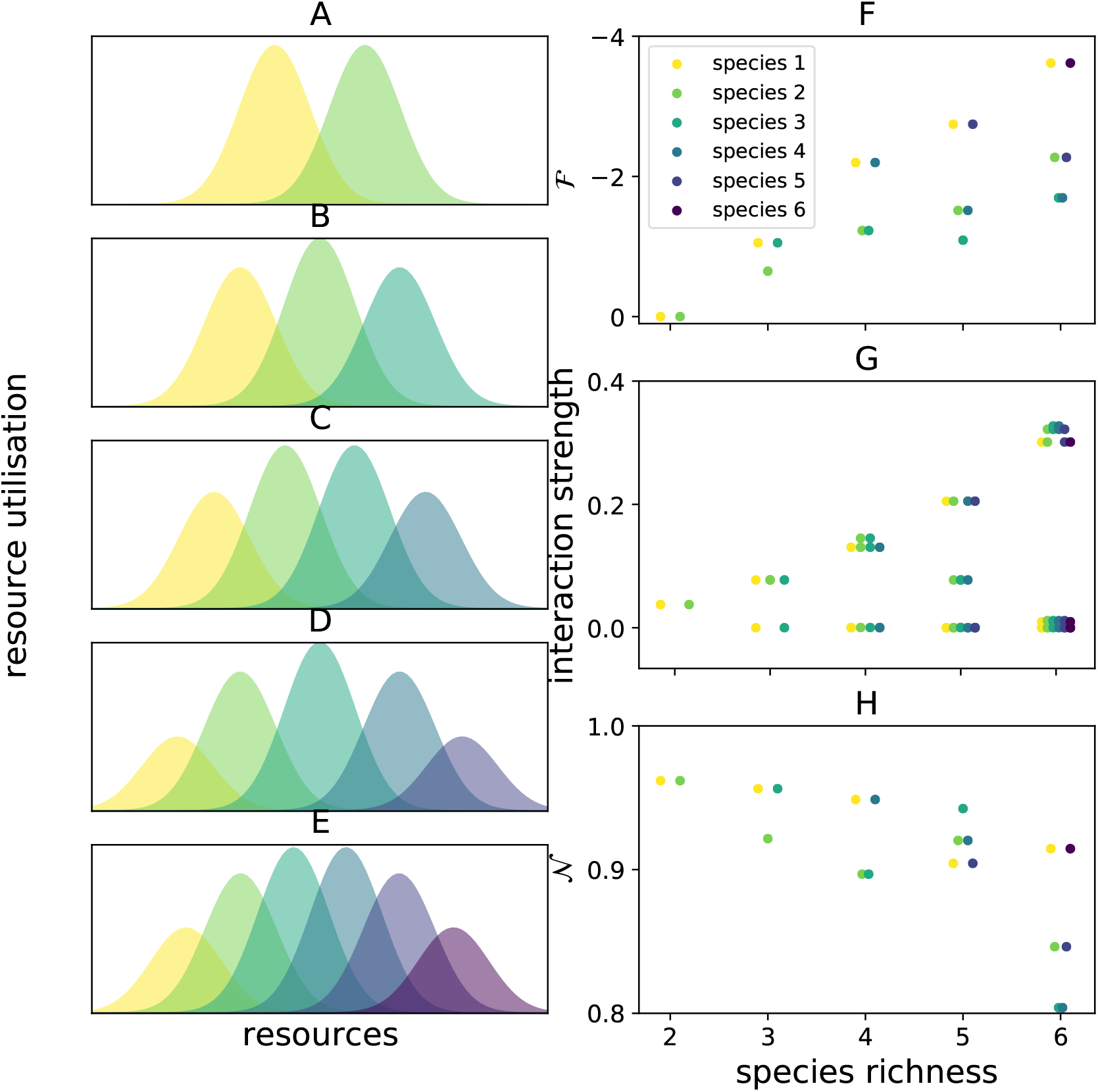
A-E: Resource utilisation of two to six competing species. Species evolve to optimal foraging strategies according to Barabás *et al*. (2016a). F: Increasing species richness increases fitness differences, as predicted by theory G: As the niche width is limited, species are located closer to each other in species rich communities and therefore have a stronger interaction strength on average. Each dot represents one value of the interaction matrix *α*_*ij*_. H: Consequently, niche differences decrease with species richness, contrary to theory. The example deviates from the theoretical predictions, as one of the key hypothesis of random assembly is not meat.

## Discussion

It is well-established that the likelihood for stable coexistence drops with species richness (Godoy *et al*., 2017; Goh & Jennings, 1977; May, 1972; Serván *et al*., 2018). Here, we explain this well known result in the context of modern coexistence theory by examining how niche and fitness differences (𝒩_*i*_ and ℱ_*i*_) change with species richness. We found that species richness, *per se*, does not affect 𝒩_*i*_ but does increase ℱ_*i*_. This conclusion is based on four independent approaches: mathematical computation, biological intuition, numerical simulations, and metanalysis of experimental data. Overall, the influence of species richness on 𝒩_*i*_ and ℱ_*i*_ is robust to inclusion or omission of the complexities (1)-(4), and all their combinations. The fitness differences of a species increases with species richness, as fitness differences measure the fitness of a species compared to the combined fitness of all other species. In multi-species communities, most species will therefore have negative fitness differences, as rarely one species will have higher fitness than all other species combined.

The niche differences of a species measure the proportion of limiting factors, e.g. resources, that are limiting to other species as well. Increasing species richness increases the amount of limiting factors shared with other species, but also the amount of limiting factors that are not shared with other species. The proportion of shared limiting factors is therefore unaffected, on average. Species-rich communities are therefore less likely to coexist (all else being equal), as fitness differences become to strong to be overcome by niche differences.

### Limitations

The available experimental data only represented fully connected communities, with no correlation among interactions (complexity (2)) and, most notably, did not contain cases of higher-order interactions (complexity (3)). We do therefore not know whether the parameter values used to describe these higher-order interactions in our simulations (and therefore the simulation results) are realistic. The available experimental data were biased towards competitive communities of terrestrial plants with relatively low species richness. Our simulations suggest that our conclusions hold for other networks as well, but we were not able to support this claim with empirical data. Computing 𝒩_*i*_ and ℱ_*i*_ on a larger collection of natural communities would help to refine our understanding of this process. However, obtaining the full interaction matrix for species-rich communities is challenging. To obtain interaction matrices, various approaches exist. For example, one uses the frequency of interaction between species (e.g. number of visits of a pollinator on a plant) as a proxy for interaction strength. The robustness of this approach, however, still needs to be tested (García-Callejas *et al*., 2018). Other methods consist of estimating interaction strength based on, for example, biomass (Moore *et al*., 1996; Zhao *et al*., 2019), mass ratio (Emmerson & Raffaelli, 2004) or production and consumption rates of species (Christensen V. & D., 1992; Jacquet *et al*., 2016). These different methods rely on different assumptions and may therefore influence the resulting matrix estimate (Carrara *et al*., 2015).

Given these limitations, one can ask to what extent addressing them would change our conclusions. In communities where species richness increases total abundance, which is the case for various communities (Grace *et al*., 2016; Loreau, 2004; Turnbull *et al*., 2013), we expect the no-niche growth rate 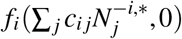 to become more negative, as 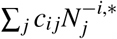 increases (eq. 2, 3). Consequently, we expect species richness to increase fitness differences, i.e. make ℱ_*i*_ more negative, as it is linear in the no-niche growth rate. Conversely, in communities where species richness decreases total abundance we expect the opposite, that is: fitness differences might decrease with species richness. It is less clear how species richness will affect niche differences in models not explored in the current paper, e.g. with different per-capita growth rates functions *f*_*i*_, or with a community with age structure (Chu & Adler, 2015). Niche differences depend on the invasion growth rate and the no-niche growth rates, which both depend on the species richness and total abundance. When species richness has a stronger negative effect on the no-niche growth rate than on the invasion growth rate, then niche differences will increase with species richness. If the invasion growth rate is decreases more, niche differences will decrease.

Additionally, we have mostly took the simplifying assumption that species richness is independent of other factors such as the average interaction strength, connectance or correlation (Cohen & Briand, 1984; Kokkoris *et al*., 2002).

The main determinant of niche differences is the average interaction strength, which might decrease with species richness due to coexistence requirements (Fig. 2A) or might increase due to increasing overlap of resource requirements (Fig. 5 G).

### New insights

Our results yield two new insights, other than the main result on how 𝒩_*i*_ and ℱ_*i*_ varies with species richness.

A first insight is that negative niche differences do not necessarily preclude coexistence. Negative niche differences have been attributed to priority effects and therefore viewed as precluding coexistence (Fukami *et al*., 2016; Ke & Letten, 2018). Our framework confirms this finding for the case of competitive two-species communities (Spaak & De Laender, 2020). However, in contrast to species in two-species communities, species in multi-species communities will not all have the same niche differences (example community in Fig 4). This implies that species *a*, with negative niche differences and low fitness differences, can coexist with species *b* and *c* that have positive niche differences and negative fitness differences. Consequently, multiple species can have negative niche differences in a multi-species communities and still persist. In our empirical data set, we found six three-species communities in which all but one species had negative niche differences. In general, we argue that a community in which all species have negative niche differences and coexist is theoretically possible. However, the kind of model and how it should be parametrized remains to be examined.

A second insight is that one can infer 𝒩_*i*_ and ℱ_*i*_ in multi-species communities from 𝒩_*i*_ and ℱ_*i*_ measured in pairwise interaction experiments. If one measures 𝒩_*i*_ and ℱ_*i*_ for each two-species sub-community of an *n* species community, which is typically done (Gallego *et al*., 2019; Godoy & Levine, 2014; Narwani *et al*., 2013; Petry *et al*., 2018), one can estimate 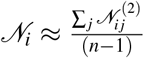. With one additional multi-species experiment to estimate the relative yield RY_*i*_ we obtain an estimation of 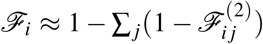. RY_*j*_ as well. This indicates that two-species experiments are sufficient to predict 𝒩_*i*_ and ℱ_*i*_ in multi-species communities.

Finally, one of the key questions in community ecology is whether niche differences are strong enough to overcome fitness differences and allow coexistence. Often, niche differences are found to be not only sufficiently strong, but much stronger than strictly needed (Chu & Adler, 2015; Levine & HilleRisLambers, 2009). The present results offer a potential explanation for this observation. That is, niche differences not only need to be sufficiently strong to overcome fitness differences of one or few competitors, as typically considered in empirical studies, but sufficiently strong to overcome fitness differences of the entire resident community, as niche differences is independent of species richness. Our results therefore allow asking the more general question of how many species one can pack in a community, given its niche difference.

## Acknowledgements

F.D.L. received support from grants of the University of Namur (FSR Impulsionnel 48454E1) and the Fund for Scientific Research, FNRS (PDR T.0048.16).

