## Appendix for "Species richness increases fitness differences, but does not affect niche differences"

### Supporting Information

### S1 Definition of niche and fitness differences

#### S1.1 Two-species communities

For simplicity we first focus on a two-species community, where we want to compute niche and fitness differences of the focal species $i$ competing with species $j$. To compute niche and fitness differences we first have to compute the conversion factors $c_{ij}$. Conceptually, the conversion factors convert density of species $j$ to species $i$ and vice versa, we therefore have

$$c_{ij}c_{ji} = 1 \tag{S1}$$

as converting species $j$ to species $i$ and back should not change the density of species $j$.

Additionally, the conversion factors convert densities of species $j$ to densities of species $i$, while leaving total density constant. Two communities with densities $(0, N_j^*)$ and $(c_{ij}N_j^*, 0)$ therefore differ only in their frequencies of species $i$ and $j$, but not in their total densities.

In a two species community, there are four sensitivities of species, the sensitivity of species $i$ to both, species $i$ and species $j$, and the sensitivity of species $j$ to both, species $i$ and species $j$. We will denote $S_{ij}$ as the sensitivity of the growth rate of species $i$ to the density of species $j$, these are respectively defined as (see also Carroll *et al.* (2011))

$$S_{ii} = \left|1 - \frac{f_i(c_{ij}N_j^*, 0)}{f_i(0,0)}\right| \tag{S2}$$

$$S_{ij} = \left|1 - \frac{f_i(0, N_j^*)}{f_i(0,0)}\right| \tag{S3}$$

$$S_{ji} = \left|1 - \frac{f_j(0, N_i^*)}{f_j(0,0)}\right| \tag{S4}$$

$$S_{jj} = \left|1 - \frac{f_j(c_{ji}N_i^*, 0)}{f_j(0,0)}\right| \tag{S5}$$

We normalize the growth rates by the intrinsic growth rate $f_i(0,0)$ for scale independency. For example, the growth rate of a grass will be much faster than the growth rate of a tree, normalizing by the intrinsic growth rates of the species allows us to compare their sensitivities. We compare the growth rates to 1, to ensure that if species $i$ is not sensitive to the density of species $j$ it has zero sensitivity (e.g. $f_i(0,0) = f_i(0, N_j^*) \Rightarrow S_{ij} = 0$). We take the absolute values, as we are interested in the magnitude of the sensitivities, not the sign.

$\sqrt{S_{ii}S_{ji}}$ is the average sensitivity of species $i$ and $j$ to the presence of species $i$, i.e. the density dependence of the community with respect to the density of species $i$. We take the geometric mean of the effects, because the effects are multiplicative quantities. By definition, the conversion factors $c_{ij}$ change frequency, but not total density. Consequently, we can compute $c_{ij}$ by solving the equation

$$\sqrt{S_{ii}S_{ji}} = \sqrt{S_{ij}S_{jj}} \tag{S6}$$

We can reorganize this equation to obtain equation (10) form Spaak & De Laender (2020):

$$\sqrt{S_{ii}S_{ji}} = \sqrt{S_{ij}S_{jj}} \tag{S7}$$

$$\left|1 - \frac{f_i(c_{ij}N_j^*, 0)}{f_i(0,0)}\right| \cdot \left|1 - \frac{f_j(0, N_i^*)}{f_j(0,0)}\right| = \left|1 - \frac{f_i(0, N_j^*)}{f_i(0,0)}\right| \cdot \left|1 - \frac{f_j(c_{ji}N_i^*, 0)}{f_j(0,0)}\right| \tag{S8}$$

$$\frac{\left|1 - \frac{f_j(0,N_i^*)}{f_j(0,0)}\right|}{\left|1 - \frac{f_j(c_{ji}N_i^*,0)}{f_j(0,0)}\right|} = \frac{\left|1 - \frac{f_i(0,N_j^*)}{f_i(0,0)}\right|}{\left|1 - \frac{f_i(c_{ij}N_j^*,0)}{f_i(0,0)}\right|} \tag{S9}$$

$$\left|\frac{f_j(0,0) - f_j(0, N_i^*)}{f_j(0,0) - f_j(c_{ji}N_i^*, 0)}\right| = \left|\frac{f_i(0,0) - f_i(0, N_j^*)}{f_i(0,0) - f_i(c_{ij}N_j^*, 0)}\right| \tag{S10}$$

Spaak & De Laender (2020) have shown that this equation always has a solution and that further more this solution is unique if we assume that $f_i$ is decreasing in $N_i$.

#### S1.2 Multi-species communities

For a multi-species communities we define niche and fitness differences equivalently to how they are defined in a two species community. The intrinsic growth rate $f_i(0, \mathbf{0})$ and the invasion growth rate $f_i(0, \mathbf{N}^{(-i,*)})$ are defined similar to the two species case, where $\mathbf{0}$ denotes the absence of all non-focal species and $\mathbf{N}^{(-i,*)}$ is the equilibrium density of the resident community. By definition, the no-niche growth rate is the growth rate of species $i$ if all other species occupied the same niche as the focal species, i.e. $f_i(\sum_j c_{ij} N_j^{(-i,*)}, \mathbf{0})$. With these three growth rates defined we can define niche and fitness differences simlar to the two species case as

$$\mathcal{N}_i = \frac{f_i(0, \mathbf{N}^{-i,*}) - f_i(\sum_{j \neq i} c_{ij} N_j^{-i,*}, \mathbf{0})}{f_i(0, \mathbf{0}) - f_i(\sum_{j \neq i} c_{ij} N_j^{-i,*}, \mathbf{0})} \tag{S11}$$

$$\mathcal{F}_i = \frac{f_i(\sum_{j \neq i} c_{ij} N_j^{-i,*}, \mathbf{0})}{f_i(0, \mathbf{0})} \tag{S12}$$

We only have to compute the conversion factors $c_{ij}$. The conversion factor $c_{ij}$ converts species $i$ and $j$, the densities of all the other non-focal species $k$ should therefore not be included into the computation of $c_{ij}$. Rather, they can be interpreted as an environmental condition. They will, however, have a potential effect on $c_{ij}$, as they may alter the availability of the limiting resources, We therefore define

$$f_i'(N_i, N_j) = f_i(N_i, N_j, \mathbf{N}_{\neg j}^{(-i,*)}) \tag{S13}$$

where $\mathbf{N}_{\neg j}^{(-i,*)}$ is the density of all other species $k$ ($k \neq i, k \neq j$) at the equilibrium density of the resident community without species $i$ present. That is $\mathbf{N}_{\neg j}^{(-i,*)})_k = N_k^{(-i,*)}$. We then solve the equations S1 and S6 from above with the function $f_i'$ and $f_j'$ to obtain $c_{ij}$ in the multi-species community.

### S2 Analytical computation of niche and fitness differences

#### S2.1 First order interactions

In this section we assume that higher order interactions are absent, i.e. $\beta_{ijk} = \gamma_{ijkl} = 0$. This case is equivalent to the Lotka-Volterra equations. This case has already been solved in the appendix of Spaak & De Laender (2020), we merely repeat their findings here. The niche difference between species $i$ and $j$ in the multispecies community is

$$\frac{f_i(0, \boldsymbol{N}^{-\boldsymbol{i},*}) - f_i(c_{ij}N_j^{-i,*}, \boldsymbol{N}_{\neg \boldsymbol{j}}^{-\boldsymbol{i},*})}{f_i(0, \boldsymbol{0}) - f_i(c_{ij}N_j^{-i,*}, \boldsymbol{N}_{\neg \boldsymbol{j}}^{-\boldsymbol{i},*})} = \tag{S14}$$

$$= \frac{\left(1 - \sum_{k\neq i} \alpha_{ik}N_k^{-i,*}\right) - \left(1 - \sum_{k\neq i,j} \alpha_{ik}N_k^{-i,*} - \alpha_{ii}c_{ij}N_j^{-i,*}\right)}{\left(1 - \sum_{k\neq i,j} \alpha_{ik}N_k^{-i,*}\right) - \left(1 - \sum_{k\neq i,j} \alpha_{ik}N_k^{-i,*} - \alpha_{ii}c_{ij}N_j^{-i,*}\right)} \tag{S15}$$

$$= \frac{\left(c_{ij}\alpha_{ii} - \alpha_{ij}\right) N_j^{-i,*}}{c_{ij}\alpha_{ii}N_j^{-i,*}} \tag{S16}$$

$$= 1 - \frac{\alpha_{ij}}{c_{ij}\alpha_{ii}} \tag{S17}$$

The absence of species $j$ in $\boldsymbol{N}_{\neg \boldsymbol{j}}^{-\boldsymbol{i},*}$ is handled by omitting this index in the summation. By solving $|1 - \frac{\alpha_{ij}}{c_{ij}\alpha_{ii}}| = |1 - \frac{\alpha_{ji}}{c_{ij}^{-1}\alpha_{jj}}|$ we get $c_{ij} = \sqrt{\left|\frac{\alpha_{jj}\alpha_{ij}}{\alpha_{ii}\alpha_{ji}}\right|}$. $c_{ij}$ is set to 0 if this value is not defined (i.e. $\alpha_{ji} = 0$) (Spaak & De Laender, 2020). For a two species community we therefore get

$$1 - \mathcal{N}_{ij}^{(2)} = sign(a_{ij})\sqrt{\left|\frac{\alpha_{ij}\alpha_{ji}}{\alpha_{ii}\alpha_{jj}}\right|} \tag{S18}$$

$$1 - \mathcal{F}_{ij}^{(2)} = \sqrt{\left|\frac{\alpha_{ii}\alpha_{ij}}{\alpha_{jj}\alpha ji}\right|} \tag{S19}$$

With this we can show that $1 - \mathcal{N}_i$ is a weighted average and $1 - \mathcal{F}_i$ is a weighted sum:

$$1 - \mathcal{N}_i = \frac{1 - \left(1 - \sum_{j \neq i} \alpha_{ij} N_j^{-i,*}\right)}{1 - \left(1 - \sum_{j \neq i} c_{ij} \alpha_{ii} N_j^{-i,*}\right)} \tag{S20}$$

$$= \frac{\sum_{j \neq i} \frac{\alpha_{ij}}{a_{ii}} N_j^{-i,*}}{\sum_{j \neq i} c_{ij} N_j^{-i,*}} \tag{S21}$$

$$= \frac{\sum_{j \neq i} \left(1 - \mathcal{N}_{ij}^{(2)}\right) c_{ij} N_j^{-i,*}}{\sum_{j \neq i} c_{ij} N_j^{-i,*}} \tag{S22}$$

$$1 - \mathcal{F}_i = 1 - \left(1 - \sum_{j \neq i} c_{ij} \alpha_{ii} N_j^{-i^*}\right) \tag{S23}$$

$$= \sum_{j \neq i} \sqrt{\left| \frac{\alpha_{ij} \alpha_{ii}}{\alpha_{ji} \alpha_{jj}} \right|} \alpha_{jj} N_j^{-i,*} \tag{S24}$$

$$= \sum_{j \neq i} (1 - \mathcal{F}_{ij}^{(2)}) \frac{N_j^{-i,*}}{N_j^*} \tag{S25}$$

Which proves the equations 4 and 5 in the main-text.

To obtain the case where $\alpha_{ij} = \bar{\alpha}$ we remark that $c_{ij} = 1$, $\mathcal{N}_{ij}^{(2)} = \bar{\alpha}$,

$\mathcal{F}^{(2)}_{ij} = 0$, $N_j^{-i,*} = \frac{1}{1+(n-2)\bar{\alpha}}$ and $N_j^*$. This leads to

$$\mathcal{N}_i = 1 - \frac{\sum_{j\neq i}(1-\mathcal{N}^{(2)}_{ij})c_{ij}N_j^{-i,*}}{\sum_{j\neq i} c_{ij}N_j^{-i,*}} \tag{S26}$$

$$= 1 - \frac{\sum_{j\neq i}(1-\bar{\alpha})\frac{1}{1+(n-2)\bar{\alpha}}}{\sum_{j\neq i}\frac{1}{1+(n-2)\bar{\alpha}}} \tag{S27}$$

$$= 1 - \bar{\alpha} \tag{S28}$$

$$\mathcal{F}_i = 1 - \sum_{j\neq i}(1-\mathcal{F}^{(2)}_{ij})\frac{N_j^{-i,*}}{N_j^*} \tag{S29}$$

$$= 1 - \sum_{j\neq i}(1-0)\frac{\frac{1}{1+(n-2)\bar{\alpha}}}{1} \tag{S30}$$

$$= 1 - \frac{n-1}{1+(n-2)\bar{\alpha}} \tag{S31}$$

#### S2.2 Higher order interactions

We assume $\alpha_{ij} = \bar{\alpha}$ and $\beta_{ijk} = \bar{\beta}$ and omit the overbars in this subsection for notational simplicity. By symmetry we have $N_j^{-i,*} = N_k^{-i,*} = N^{-i,*}$ and $c_{ij} = c_{ji} = 1$.

$$\mathcal{N}_i = \frac{r_i\left(1-\sum_j \alpha N^{-i,*}\left(1+\sum_k \beta N^{-i,*}\right)\right) - r_i\left(1-\sum_j N^{-i,*}\left(1+\beta\sum_k N^{-i,*}\right)\right)}{r_i - r_i\left(1-\sum_j N^{-i,*}\left(1+\beta\sum_k N^{-i,*}\right)\right)} \tag{S32}$$

$$= \frac{(1-\alpha)(n-1)N^{-i,*}(1+\beta(n-1)N^{-i,*})}{(n-1)N^{-i,*}(1+\beta(n-1)N^{-i,*})} \tag{S33}$$

$$= 1-\alpha \tag{S34}$$

To compute $\mathcal{F}_i$ we first make some observations about the equilibrium density $N^{-i,*}$:

$$0 = 1 - \left(N^{-i,*} + \alpha(n-2)N^{-i,*}\right)\left(1 + \beta(n-1)N^{-i,*}\right) \tag{S35}$$

$$1 = \left(N^{-i,*} + \alpha(n-2)N^{-i,*}\right)\left(1 + \beta(n-1)N^{-i,*}\right) \tag{S36}$$

$$\frac{1}{N^{-i,*} + \alpha(n-2)N^{-i,*}} = 1 + \beta(n-1)N^{-i,*} \tag{S37}$$

$$\mathcal{F}_i = 1 - (n-1)N^{-i,*}(1 + \beta(n-1)N^{-i,*}) \tag{S38}$$

$$= 1 - \frac{(n-1)N^{-i,*}}{N^{-i,*} + \alpha(n-2)N^{-i,*}} \tag{S39}$$

$$= 1 - \frac{n-1}{1-(n-2)\alpha} \tag{S40}$$

Higher order interactions therefore do not affect $\mathcal{N}$ and $\mathcal{F}$ on average.

#### S2.3 Indirect effects

To investigate complexity (4) we removed indirect effects. Indirect effects are defined as a third species $k$ affecting densities of the non-focal species $j$, which affects the effect of species $j$ on the focal species $i$. To remove these indirect effects we set $N_j^{-i,*} = N_j^*$, i.e. a species $k$ does not affect the density of species $j$. Note however, that species $k$ can still affect species $i$ directly via $\alpha_{ik}$ or via higher-order effects (e.g. $\beta_{ijk}$).

$$\mathcal{F}_i = 1 - \sum_{\alpha_{ij}\neq 0} (1 - \mathcal{F}_{ij}^{(2)}) \frac{N_j^{-i,*}}{N_j^*} \tag{S41}$$

$$= 1 - \sum_{\alpha_{ij}\neq 0} (1 - \mathcal{F}_{ij}^{(2)}) \frac{N_j^*}{N_j^*} \tag{S42}$$

$$= 1 - \sum_{\alpha_{ij}\neq 0} (1 - \mathcal{F}_{ij}^{(2)}) \tag{S43}$$

$$\mathcal{N}_i = \frac{\sum_{\alpha_{ij}\neq 0}(1 - \mathcal{N}_{ij}^{(2)}) N_j^{-i,*}}{\sum_{\alpha_{ij}\neq 0} c_{ij} N_j^{-i,*}} \tag{S44}$$

$$= \frac{\sum_{\alpha_{ij}\neq 0}(1 - \mathcal{N}_{ij}^{(2)}) c_{ij} N_j^*}{\sum_{\alpha_{ij}\neq 0} c_{ij} N_j^*} \tag{S45}$$

$\mathcal{F}_i$ changes from a saturating to a linear response in species richness, i.e. $\mathcal{F}_i \approx 1 - (n-1)$ on average. Conversely, removing indirect effects will not change $\mathcal{N}_i$ on average. Thus, indirect effects will mostly not change the response of $\mathcal{N}_i$ to species richness. This yields an important result: Indirect effects are purely equalizing as they do not change niche differences, and thus promote coexistence.

### S3 Simulations

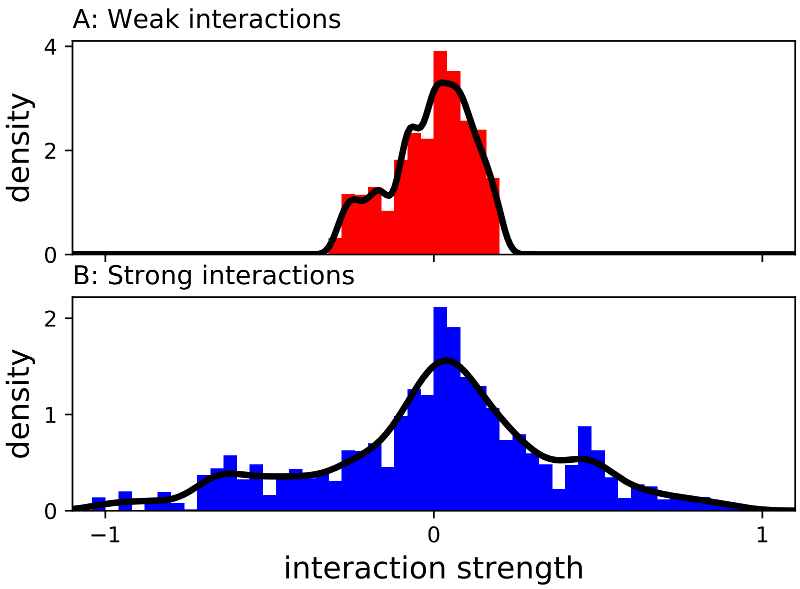


**Figure S1:** Interaction strength distributions for weak (A) and strong (B) first order interactions. Histogram shows the distribution of the interaction strength of empirical communities that coexist. We fit a guassian kernel density distribution (black line) to the distribution of the interaction strength of empicial communities that coexist (histogram). We removed outliers, defined as interaction strength that were below $Q_1 - 1.5(Q_3 - Q_1)$ or above $Q_3 + 1.5(Q_3 - Q_1)$, where $Q_1$ and $Q_3$ are the first and third quartile. For weak interaction strength we only retained interaction strength within the first and third quartile.

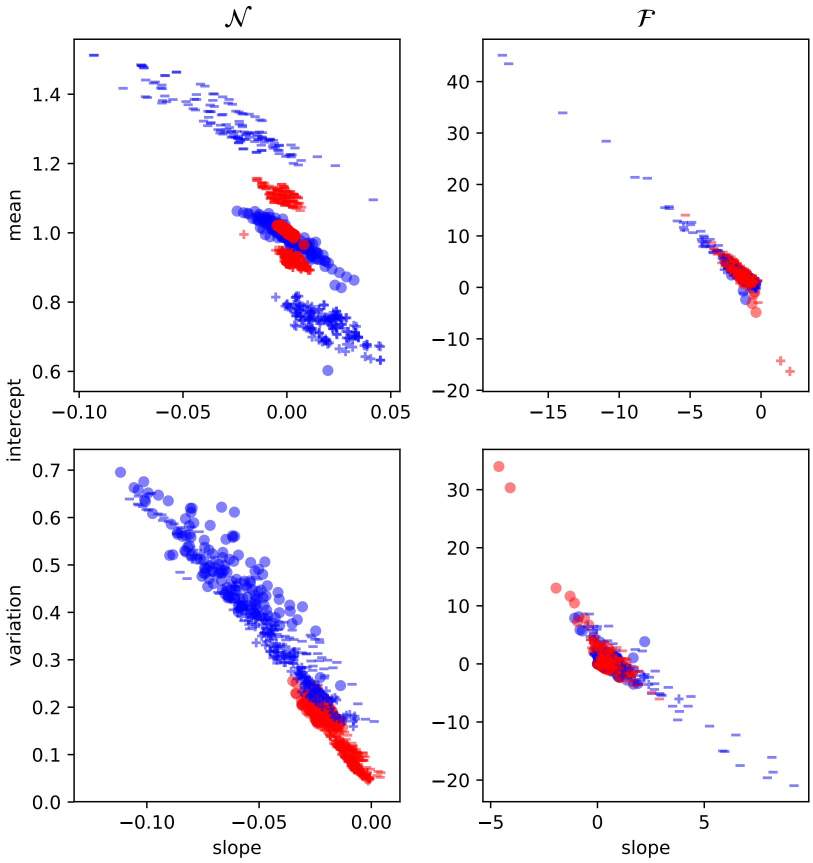


**Figure S2:** Effects of species richness on $\mathcal{N}$ and $\mathcal{F}$ (top row) and on variation of $\mathcal{N}$ and $\mathcal{F}$ (bottom row) per factor combination of the full-factorial design. For each factor combination we fit a linear regression of the response variable as a function of species richness. X-axis correspond to the slope, y-axis correspond to the intercept of the linear regression. Color codes the strength of first order interaction: red: weak, blue: strong. Shape codes the type of first order interactions: minus: facilitation, plus: competition, dot: both. The first order interaction affects the intercept of $\mathcal{N}$ and variation of $\mathcal{N}$ most. Variation of $\mathcal{N}$ decreased for almost all factor combinations. $\mathcal{F}$ decreased in almost all factor combination.

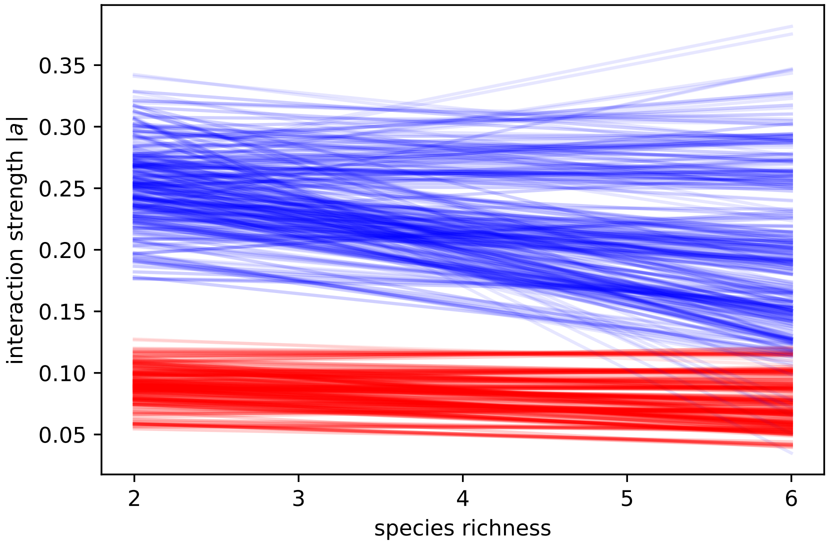


**Figure S3:** Effects of species richness on absolute interaction strength. The y-axis shows the mean of the absolute interspecific-interaction strength. Each line is a linear fit of species richness versus interaction strength in one of the full-factorial settings. Species richness affects interaction strength in communities with strong first order interactions (blue lines) mostly negatively. Species richness has a minor effect on interaction strength in communities with weak first order interactions (red lines).

| | ND slope | ND var slope | FD slope | FD var slope |
|---|---|---|---|---|
| ord1: negative | [-0.014; 0.003] | [-0.027; -0.005] | [-3.003; -0.915] | [0.001; 1.446] |
| ord1: unrestricted | [-0.004; 0.003] | [-0.031; -0.016] | [-1.808; -0.607] | [-0.43; 0.772] |
| ord1: positive | [-0.002; 0.009] | [-0.016; -0.002] | [-1.978; -0.577] | [-0.125; 0.72] |
| ord2: negative | [-0.014; 0.008] | [-0.029; -0.004] | [-2.11; -0.817] | [-0.05; 0.889] |
| ord2: absent | [-0.01; 0.008] | [-0.029; -0.004] | [-2.379; -0.707] | [-0.091; 1.438] |
| ord2: positive | [-0.014; 0.007] | [-0.029; -0.007] | [-2.837; -0.746] | [-0.942; 1.276] |
| ord2: unrestricted | [-0.005; 0.003] | [-0.025; -0.002] | [-2.192; -0.564] | [0.02; 0.921] |
| ord3: absent | [-0.01; 0.008] | [-0.026; -0.005] | [-2.688; -0.588] | [-0.199; 1.133] |
| ord3: present | [-0.013; 0.007] | [-0.029; -0.003] | [-2.532; -0.692] | [0.097; 1.253] |
| con: high | [-0.014; 0.009] | [-0.029; -0.005] | [-2.521; -0.563] | [-0.113; 1.251] |
| con: middle | [-0.006; 0.003] | [-0.027; -0.002] | [-2.929; -0.709] | [0.06; 1.32] |
| con: low | [-0.008; 0.005] | [-0.027; -0.005] | [-2.421; -0.751] | [-0.257; 0.965] |
| cor: negative | [-0.009; 0.006] | [-0.029; -0.001] | [-2.791; -0.737] | [0.077; 1.25] |
| cor: positive | [-0.011; 0.008] | [-0.025; -0.005] | [-2.659; -0.73] | [-0.316; 1.38] |
| cor: none | [-0.014; 0.009] | [-0.029; -0.009] | [-2.218; -0.588] | [-0.164; 1.001] |
| indirect: absent | [-0.01; 0.008] | [-0.028; -0.004] | [-2.742; -0.633] | [-0.036; 1.323] |
| indirect: present | [-0.01; 0.007] | [-0.028; -0.004] | [-2.325; -0.627] | [-0.108; 1.009] |

**Table S1:** For each factor level (row names) we show the 5% and the 95% percentiles for the slopes of $\mathcal{N}_i$, $\mathcal{F}_i$ and their interquartile range.

### S4 Literature Data

| | 2 | 3 | 4 | 5 | 6 | 7 | 8 | 9 | Total |
|---|---|---|---|---|---|---|---|---|---|
| Original matrices | 0 | 8 | 13 | 1 | 5 | 3 | 0 | 3 | 33 |
| Subcommunities | 358 | 527 | 576 | 472 | 278 | 111 | 27 | 3 | 2544 |
| int. matrix" | 356 | 517 | 557 | 455 | 271 | 110 | 27 | 3 | 2296 |
| NFD computed | 356 | 371 | 140 | 45 | 8 | 0 | 0 | 0 | 920 |
| coexistence | 290 | 270 | 142 | 55 | 11 | 0 | 0 | 0 | 768 |
| comp. exclusion | 66 | 247 | 415 | 400 | 260 | 110 | 27 | 3 | 1528 |
| no invasion analysis | 0 | 8 | 23 | 12 | 3 | 0 | 0 | 0 | 46 |
| invasion wrong | 0 | 3 | 0 | 0 | 0 | 0 | 0 | 0 | 3 |
| NFD coexistence | 290 | 262 | 119 | 43 | 8 | 0 | 0 | 0 | 722 |
| NFD comp. excl | 66 | 109 | 21 | 2 | 0 | 0 | 0 | 0 | 198 |

**Table S2:** Number of communities per species richness (top row). *Original matrices*: Number of different communities found in the literature. *Subcommunities*: Number of subcommunities generated out of all matrices. *int. matrix*: Number of communities in which all interaction strength are present. *NFD computed*: Number of communities for which we were able to compute $\mathcal{N}$ and $\mathcal{F}$. *coexistence*: Number of communities with a stable point equilibrium. *comp. exclustion*: Number of communities with no stable point equilibrium. *no invasion analysis*: Number of communities in which at least one subcommunity did not have a stable equilibrium, preventing invasion analysis. *invasion wrong*: Number of communities in which invasion as predicted by invasion analysis did not agree with coexistence as defined by stable point equilibrium. All three communities stem from the same original matrix. The species facilitate each other, such that invasion is positive, however densities explode to infinity. *NFD coexistence*: Number of communities in which $\mathcal{N}$ and $\mathcal{F}$ predict coexistence. *NFD comp. excl*: Number of communities in which $\mathcal{N}$ and $\mathcal{F}$ predict competitive exclusion.

### S5 Non random species arrival

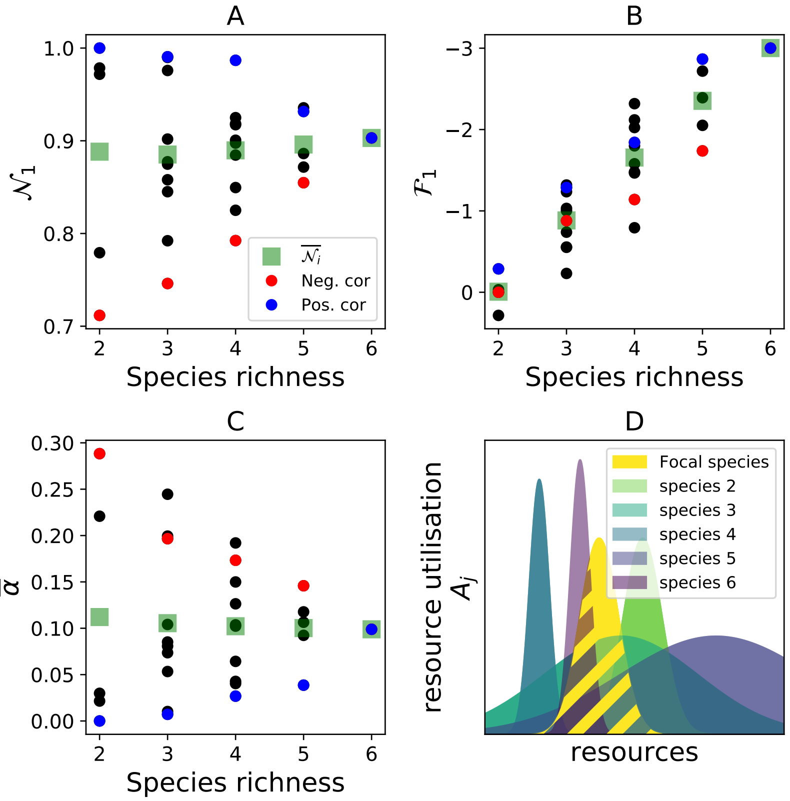


**Figure S4:** In the main text we assumed that species arrive in the usual order (2-6). In this order species richness and average interaction strength with species 1 are not correlated, i.e. random, as assumed by the theroy. Alternatively, we can order the species to arrive such that interaction strenght and species richness are positively (blue dots) or negatively (red dots) correlated. In this case, $\mathcal{N}_1$ will decrease or increase, respectively, with species richness (Panel A). However, the average of $\mathcal{N}_1$ over all communities with species 1 present is not expected to change (green squares). Note, there is only one community with 6 species present, the entire community. Panel B: Regardless of the community assembly process, fitness differences increases with species richness. Panel C: The change of $\mathcal{N}_i$ with species richness is caused by the underlying change of $\bar{\alpha}$ with species richness. Panel D: Equivalent to figure 1E from the main text.

### S6 Species packing

The species compete for resources and can evolve their resource uptake traits as proposed in Barabás *et al.* (2016). Specifically, the growth rate of species $i$ is given by

$$\frac{dN_i(t)}{dt} = N_i(t)\left(b_i(t) - \sum_j \alpha_{ij}(t)N_j(t)\right) \tag{S46}$$

,where $N_i(t)$ is the density of species $i$ at time $t$, $b_i(t)$ is the intrinsic growth rate of species $i$ at time $t$ and $\alpha_{ij}(t)$ is the per capita effect of species $j$ on species $i$ at time $t$. $b_i$ and $\alpha_{ij}$ are both time dependent, because the species may alter their resource consumption traits. They depend on the resource consumption traits of the species as follows

$$b_i = \int \exp(-\frac{(z-\mu_i(t))^2}{2\sigma^2})dz - m_i(t) \tag{S47}$$

$$\alpha_{ij} = \int \exp(-\frac{(z-\mu_i(t))^2}{2\sigma^2})\exp(-\frac{(z-\mu_j(t))^2}{2\sigma^2}) \tag{S48}$$

, where $\sigma$ is the variance of the resource consumption (assumed to be species independent), $\mu_i$ is the mean of the resource consumption of species $i$ and $m_i(t)$ is the mortality rate of species $i$, which is assumed to depend on the trait value $\mu_i$: $m_i = -\mu_i^2$, that is we assume an optimal value for $\mu_i$ and deviations from this optimum lead to a higher mortality rate. Species can evolve their trait value $\mu_i$ over time, shown in graph 4 are the endpoints of this evolution, where no species may alter their $\mu_i$ to increase their growth rate, that is $\frac{d}{d\mu_i}\left(\frac{dN_i}{dt}\right) = 0$.
